# Distinct types of multicellular aggregates in *Pseudomonas aeruginosa* liquid cultures

**DOI:** 10.1101/2022.05.18.492589

**Authors:** Gavin Melaugh, Vincent Martinez, Perrin Baker, Preston Hill, P. Lynne Howell, Daniel J. Wozniak, Rosalind J. Allen

**Affiliations:** SUPA, School of Physics and Astronomy, University of Edinburgh, United Kingdom; School of Engineering, University of Edinburgh, United Kingdom; Program in Molecular Medicine, Research Institute, The Hospital for Sick Children, Toronto, Ontario M5G 0A4, Canada; Departments of Microbial Infection and Immunity, Microbiology, Infectious Diseases Institute, Ohio State University, Columbus, OH 43210, USA; Department of Biochemistry, University of Toronto, Toronto, Ontario M5S 1A8, Canada; Theoretical Microbial Ecology, Institute of Microbiology, Faculty of Biological Sciences, Friedrich Schiller University Jena, Buchaer Strasse 6, 07745 Jena, Germany

## Abstract

*Pseudomonas aeruginosa* forms suspended multicellular aggregates when cultured in liquid media. Such aggregates may be important in disease, and/or as a pathway to biofilm formation. The polysaccharide Psl and extracellular DNA (eDNA) have both been implicated in aggregation, but previous results depend strongly on the experimental conditions. Here we develop a quantitative microscopy-based method for assessing changes in the size distribution of suspended aggregates over time in growing cultures. For exponentially growing cultures of *P. aeruginosa* PAO1, we find that aggregation is mediated by cell-associated Psl, rather than by either eDNA or secreted Psl. These aggregates arise *de novo* within the culture via a growth process that involves both collisions and clonal growth. They are “non-cheatable” since Psl non-producing cells do not aggregate with producers. In contrast, we find that stationary phase (overnight) cultures contain a different type of multicelullar aggregate, in which both eDNA and Psl mediate cohesion. Our findings suggest that the physical and biological properties of multicellular aggregates may be very different in early-stage vs late-stage bacterial cultures.

## INTRODUCTION

Bacteria tend to aggregate when suspended in liquid or viscoelastic media. In clinical settings, this can have devastating consequences. For example, in the cystic fibrosis lung, aggregates of the opportunistic pathogen *Pseudomonas aeruginosa* are implicated in life-long lung infections that can result in complete failure of the respiratory system (1, 2, 3). Bacterial aggregation is also relevant in wastewater treatment where pollutant-degrading bacteria form compact aggregates (or flocs) (4); here, unpredictability in floc formation is costly (5). From a microbiological perspective, aggregates can show similar properties to surface-attached biofilms, including antibiotic tolerance (6, 7, 8), and in some cases they may initiate biofilm formation (9, 10). Yet despite its importance, a clear picture of the mechanisms underlying aggregate formation under different environmental conditions remains lacking.

In this work, we investigate quantitatively aggregation of the *P. aeruginosa* lab strain PAO1 in liquid culture. We compare aggregates formed during exponential growth with those in overnight, stationary phase cultures. *P. aeruginosa* is widely used as a model organism for biofilm formation on surfaces (11, 12, 13, 14, 15), but it also forms aggregates in liquid culture (16, 17, 6, 9, 7). These aggregates have been observed, in different studies, in both exponential and stationary phase cultures (17, 9, 7, 16, 6), with most reports focusing on late-log or stationary phase (17, 9, 16, 6). It is unclear whether differences exist between aggregates that form in different growth phases. In late stationary phase, aggregate dispersal has been reported (17) (in a similar manner to biofilm dispersal (18)).

Extracellular polymeric substances (EPS) play a central role in the multicellular behaviour of *P. aeruginosa* (19, 20, 21, 22, 23, 24). For the PAO1 strain, three polymers have been shown to be important in the formation of biofilms on surfaces: the two polysaccharides Psl (25, 26, 27, 28, 29) and Pel (27, 28), and extracellular DNA (eDNA) (16, 27, 30, 26, 31) (in other strains, alginate production is also a significant factor). The protein CdrA has also been implicated in biofilm formation through its propensity to bind to Psl and Pel (32, 33). Of the three polymers, Psl is thought to be the predominant biofilm matrix component for the PAO1 strain (34), facilitating cell-surface and cell-cell adhesion (29, 35), as well as providing mechanical stability and structural integrity to the biofilm (26). Psl is an uncharged polymer of mannose, rhamnose, and glucose (26, 36) that exists in two forms (36, 37, 38): a cell-free form that is secreted into the medium, and a cell-associated form that remains bound to the cell surface. It is not yet known which of these two forms predominates in the biofilm matrix. *P. aeruginosa* also produces a glycoside hydrolase, PslG, that is thought to play a regulatory role in Psl production (38). PslG has been shown to specifically degrade Psl and to disrupt *P. aeruginosa* biofilms (39, 40). The Pel polysaccharide, composed of galactosamine and glucosamine sugar residues, is cationic (37). Pel plays a more major role in the PA14 strain (deficient in Psl production due to mutations in the *psl* genes), where it is required for mature biofilm formation; however in PAO1 strains that are deficient in Psl production, upregulation of Pel can restore biofilm formation (41). eDNA has been found to play a structural role in young PAO1 biofilms and is also present in the matrix of established biofilms (31), although it seems to be less important for cell-cell cohesion (30). eDNA is generated in biofilms, and in late log phase liquid cultures, via lysis of a subpopulation of bacteria, in a manner that depends on quorum sensing, flagella and pili (16, 42).

Perhaps unsurprisingly, EPS has also been implicated in liquid-phase aggregation of *P. aeruginosa* cells (16, 17, 6, 7). In late-stage (late-log or stationary phase) cultures, aggregation has been found to be mediated by eDNA (16, 17), released in a quorum-sensing mediated manner (16), or by a combination of eDNA and Psl (6, 7). It is not clear whether the aggregates are primarily formed by collisions between individual planktonic cells, or via clonal growth of a few aggregated cells, although for exponential-phase aggregates there is some evidence for a collision model (7). So far only Kragh *et al* have reported on aggregation in exponential phase cultures (7), where Psl-mediated aggregates were observed. A strong dependence on inoculation conditions suggested that these exponential phase aggregates might be seeded by pre-existing aggregates in the inoculum (7). Taken together, previous work suggests that exponential phase aggregates may be distinct from those found in late-log or stationary phase, but the details of how aggregates initiate and grow remain unclear.

The involvement of EPS in the multicellular behaviour of *P. aeruginosa* raises interesting socio-evolutionary questions (43, 44, 45, 46), since secreted EPS is shared between cells and could in principle provide a benefit to non-producing “cheaters”. Previous work on surface-attached biofilms has suggested that Psl production is a “social but non-cheatable” biofilm trait (43). In other words, the benefits of Psl are shared between cells, but non-producers do not exploit these benefits. Investigating the socio-evolutionary implications of EPS in liquid phase aggregates is of great interest, and may have wider relevance since aggregate formation has been suggested as a first step towards evolution of multicellular behaviour more generally (47, 48).

Here, we develop a quantitative image-analysis method to characterise the time evolution of the aggregate-size distribution in liquid cultures of *P. aeruginosa*. We observe distinct types of aggregates in exponential-phase cultures compared to overnight stationary-phase cultures. We show that cell-associated Psl mediates aggregation in exponential-phase cultures, while eDNA plays no significant role. We find evidence that aggregates appear *de novo* in these cultures, and grow via a mixture of collisions and clonal growth. Psl in these aggregates is “non-cheatable” since Psl non-producers do not aggregate with producers. In contrast, we find a different type of aggregate in overnight stationary phase cultures. Both Psl and eDNA are involved in these stationary-phase aggregates, but eDNA alone is suffcient to mediate aggregation. Our work helps clarify the nature of aggregation in early-stage exponential cultures and highlight the fact that different aggregation mechanisms are relevant under different conditions of bacterial growth.

## RESULTS

### PAO1 forms aggregates during exponential growth

To assess the formation of *P*.*aeruginosa* aggregates during exponential growth, we inoculated liquid LB media with a low density of cells taken from an overnight culture, and took regular samples for microscopy while tracking the growth of the cultures via optical density (OD) measurements (Fig. 1(a); see Methods). We observed exponential growth in the first 5 hours after inoculation, with growth rate ∼ 1.3 h^−1^ corresponding to a doubling time of ∼ 30 min in agreement with previous studies (49) (inset to Fig. 1(a)). Microscopy images of the liquid samples, taken in a capillary tube (see Methods), showed clearly the presence of multicellular aggregates coexisting with nonaggregated cells (Figs. 1(b) to (d)). At later times, when the culture ceased to grow exponentially, the aggregates disappeared, suggesting that they had dispersed (Fig. 1(e)).

**FIG 1.**
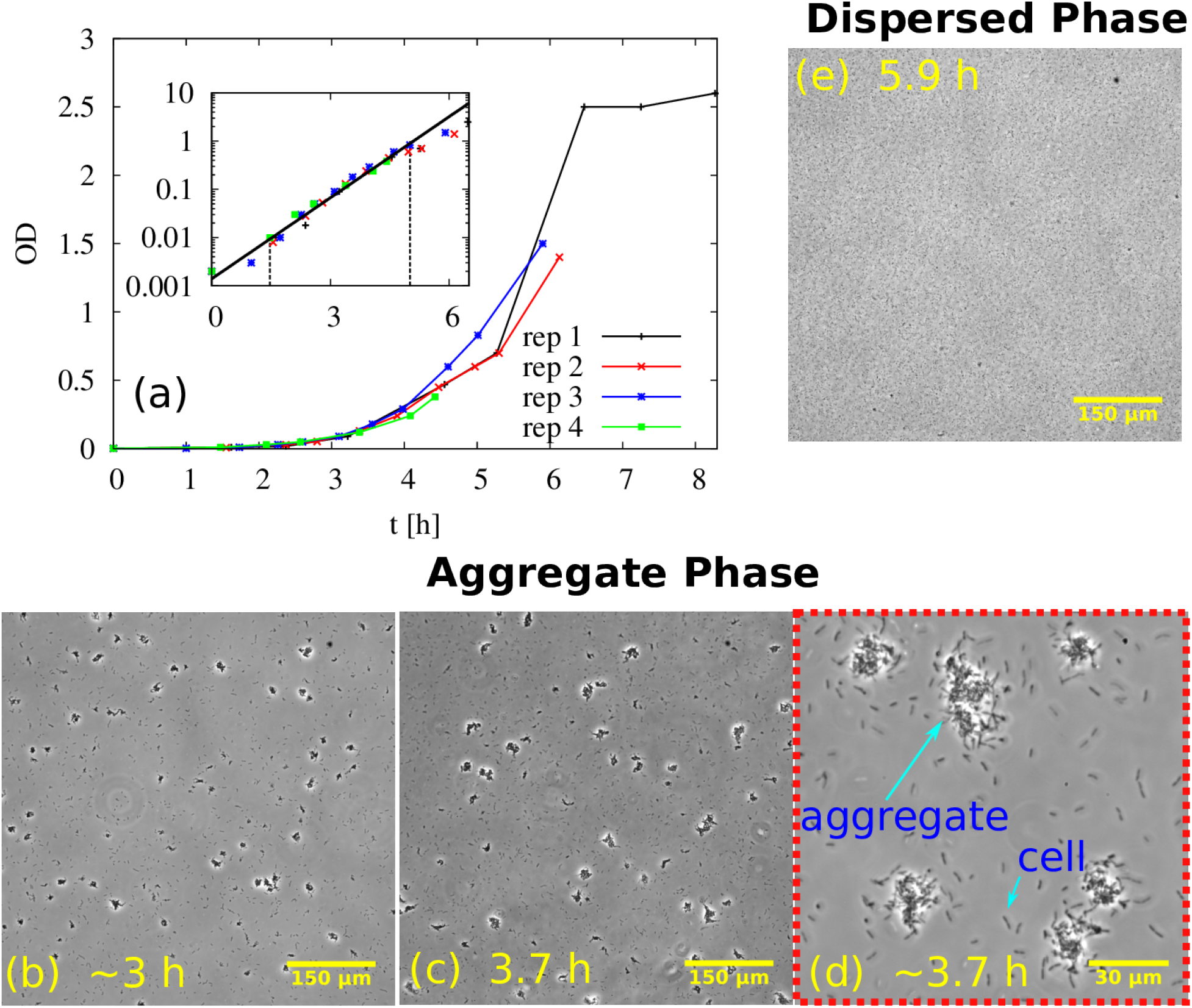
Aggregated and nonaggregated cells of PAO1 are present in the suspension during exponential growth. (a) Optical density (OD) as a function of time in 4 replicate (rep) experiments. (b)-(e) Representative phase contrast microscopy images at different times during growth. (b) 3 h, 20× magni cation. (c) 3.7 h, 20× magni cation. (d) 3.7 h, 90 × magni cation. Note the image in (d) was sampled from a different field of view to that in (c). (e) 5.9 h, 20× magni cation. The inset shows the data the 4 replicate experiments plotted on a logarithmic scale (Y axis), and line of best fit (solid black line) corresponding to a growth rate of ∼ 1.3 h^−1^ (doubling time of ∼ 30 min).

### Aggregate size distribution changes with time as aggregates grow

Visual inspection of microscopy images (Figs. 1(b) and (c)) suggests that aggregates become larger over time during exponential growth. To investigate this more quantitatively, we developed an automated image analysis protocol for measuring the cross-sectional area of individual aggregates on the bottom surface of the viewing capillary (see Methods), an approach similar to that of previous studies (50). This allowed us to obtain a probability distribution of observed aggregate sizes, for each of our samples (Fig. 2(a)). The probability *p* (*A*) of observing an aggregate of cross-sectional area *A* is a decreasing function of *A*, indicating that small aggregates are more abundant than larger ones. Nonaggregated cells were the most abundant entities in all of our samples.

**FIG 2.**
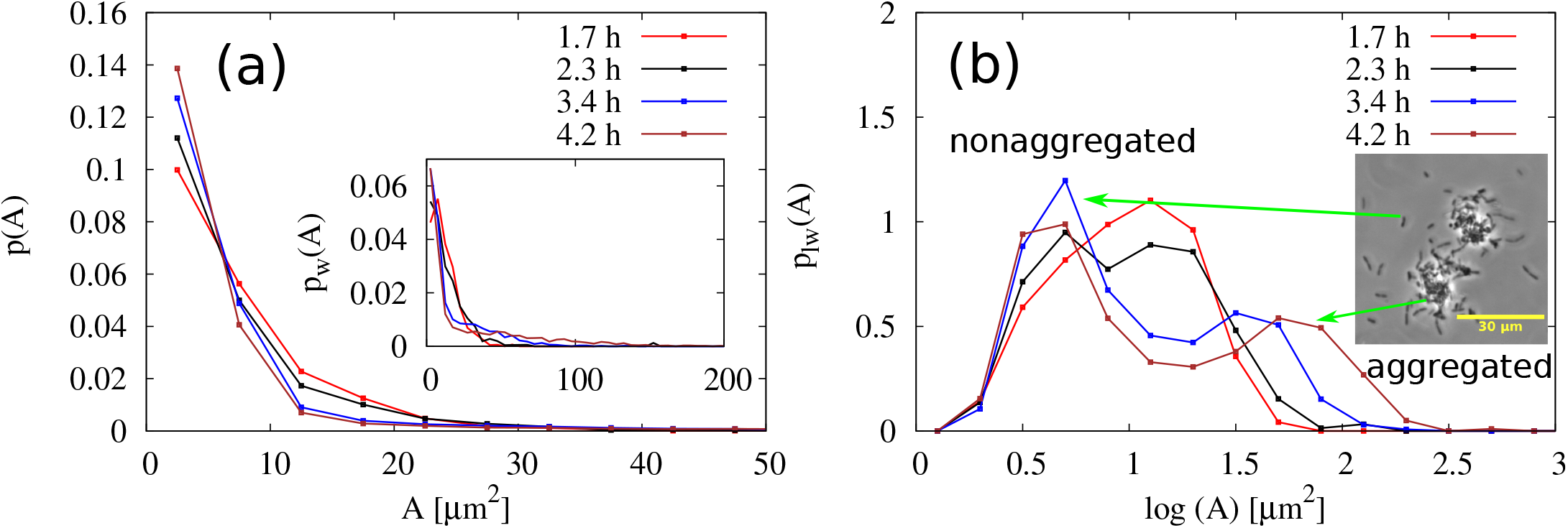
Aggregates increase in size during exponential growth. Distributions of aggregate sizes for PAO1 at 4 different times during incubation in LB. (a) The distribution of aggregate sizes. *p* (*A*) is the probability density of observing an aggregate of size *A*. Inset-weighted distribution, *p*_*w*_ (*A*) in which the frequency is multiplied by the aggregate size. (b) Logged and weighted distributions, *p*_*l w*_ (*A*), of aggregate sizes. *p*_*w*_ (*A*) is the probability density for a pixel to belong to an aggregate of size *A*. Taking the log of the distribution has the effect of compressing the x-axis which is necessary for visualisation given the broad range of aggregates in the system (see Methods). The inset shows a representative phase contrast image of an aggregate and nonaggregated cells with green arrows pointing to their inluence on the distribution.

To highlight more clearly the changes in the aggregate size distribution with time, we computed the weighted distribution *p*_*w*_ (*A*) (inset Fig. 2(a)). This highlights the contribution of larger aggregates. It consists of the raw aggregate size probability distribution *p* (*A*) (Fig. 2(a)), multiplied by the aggregate size. Hence, *p*_*w*_ (*A*) provides an estimate of the probability that a given bacterium in our sample belongs to an aggregate of size *A* (see Methods). To better visualise the contribution of the larger aggregates to the distribution, we also take the log of the aggregate size before weighting to give the log-weighted distribution *p*_*l w*_ (*A*) (Fig. 2(b)).

This form of analysis (Fig. 2(b)) clearly shows that, over time, a coexistence emerges between nonaggregated cells, to which we attribute the peak at log(*A*) ∼ 0.7 (*A* = 10^0.7^ = 5 µm^2^), and aggregates, to which we attribute the peak on the right side of the distribution log(*A*) > 1.5 (*A* > 10^1.5^ = 30 µm^2^). The “aggregate peak” shifts to the right at later times (blue and brown curve), indicating growth of the aggregates. In contrast, the nonaggregated-cell peak does not shift, consistent with our expectation that nonaggregated cells should not change in size during the exponential phase of growth.

### Both collisions and clonal growth are involved

Growth of an aggregate could happen due to proliferation of cells within the aggregate (clonal growth), or due to collisions and subsequent sticking of single cells or other aggregates from within the culture. Previous evidence for *P. aeruginosa* (7) and for *Staphylococcus aureus* (51) have suggested that both mechanisms might be involved. To assess the relative importance of these two mechanisms in the growth of our exponential phase aggregates, we repeated our experiment, this time inoculating with a 1:1 ratio (see Methods) of mCherry-labelled (red) and GFP-labelled (green) PAO1 cells (both from overnight cultures). If aggregation due solely to clonal growth (originating from a single cell) predominates, we would expect aggregates to be composed of cells of a single colour (reflecting the colour of the progenitor cell), while if growth by collisions predominates, we would expect to see a mix of colours within each aggregate. Using epifluorescence microscopy to image the aggregates that formed, we observed both red and green bacteria within the same aggregate (Fig. 3), suggesting that collision processes are indeed involved in aggregate growth. However, upon closer inspection, we found that the colour distribution within the aggregate was not entirely well mixed; instead, it consisted of red domains and green domains. This suggests that clonal growth is also occurring in the suspended exponential phase aggregates.

**FIG 3.**
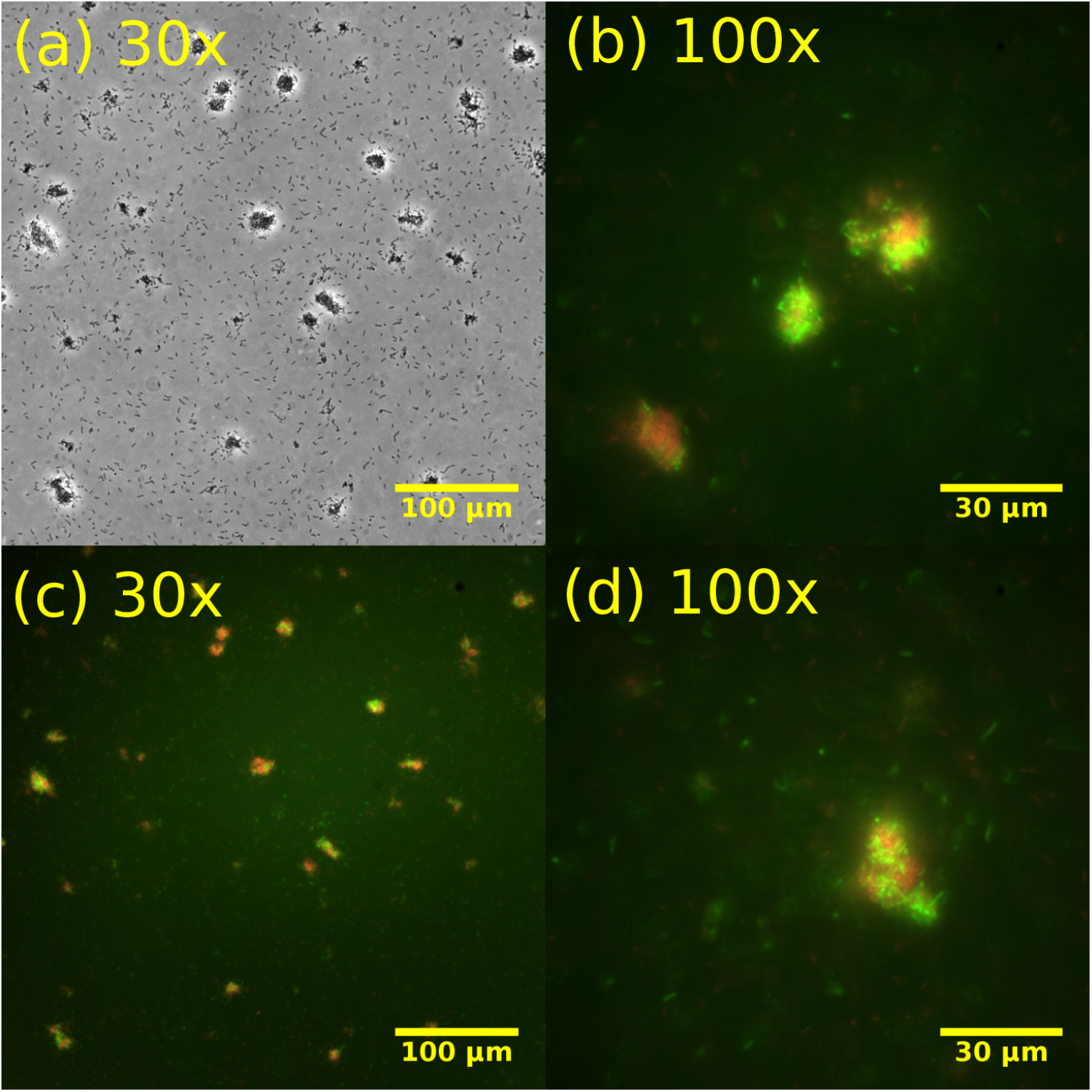
Cell-cell collisions are involved in aggregate growth. Phase contrast and luorescence microscopy images of aggregated samples of a mixed system of PAO1 GFP and PAO1 mCherry. (a) Phase contrast image at 4.8 h. (b) Superposed fluorescence images from green (PAO1 GFP) and red channels (PAO1 mCherry) coresponding to (a). (c) and (d) Superposed fluorescence images from green (PAO1 GFP) and red channels (PAO1 mCherry) at 5.0 h.

### Psl is responsible for exponential phase aggregation

To assess the involvement of the polysaccharides Pel and Psl in aggregate formation, we used three *P. aeruginosa* strains with different deficiencies in polysaccharide production: the mutant Δ*pel*, which can produce Psl but not Pel; the mutant Δ*psl*, which can produce Pel but not Psl, and the double mutant Δ*pel*Δ*psl*, which cannot produce either Psl or Pel. We observed no aggregates in samples taken from exponential cultures of the two non-Psl producing strains Δ*psl* and Δ*pel*Δ*psl*; this was also reflected in the lack of an aggregate signal in the weighted size distribution *p*_*l w*_ (*A*) (Figs. 4(a)-(d)). Therefore, these data suggest that Psl is essential for aggregate formation in our experiments.

**FIG 4.**
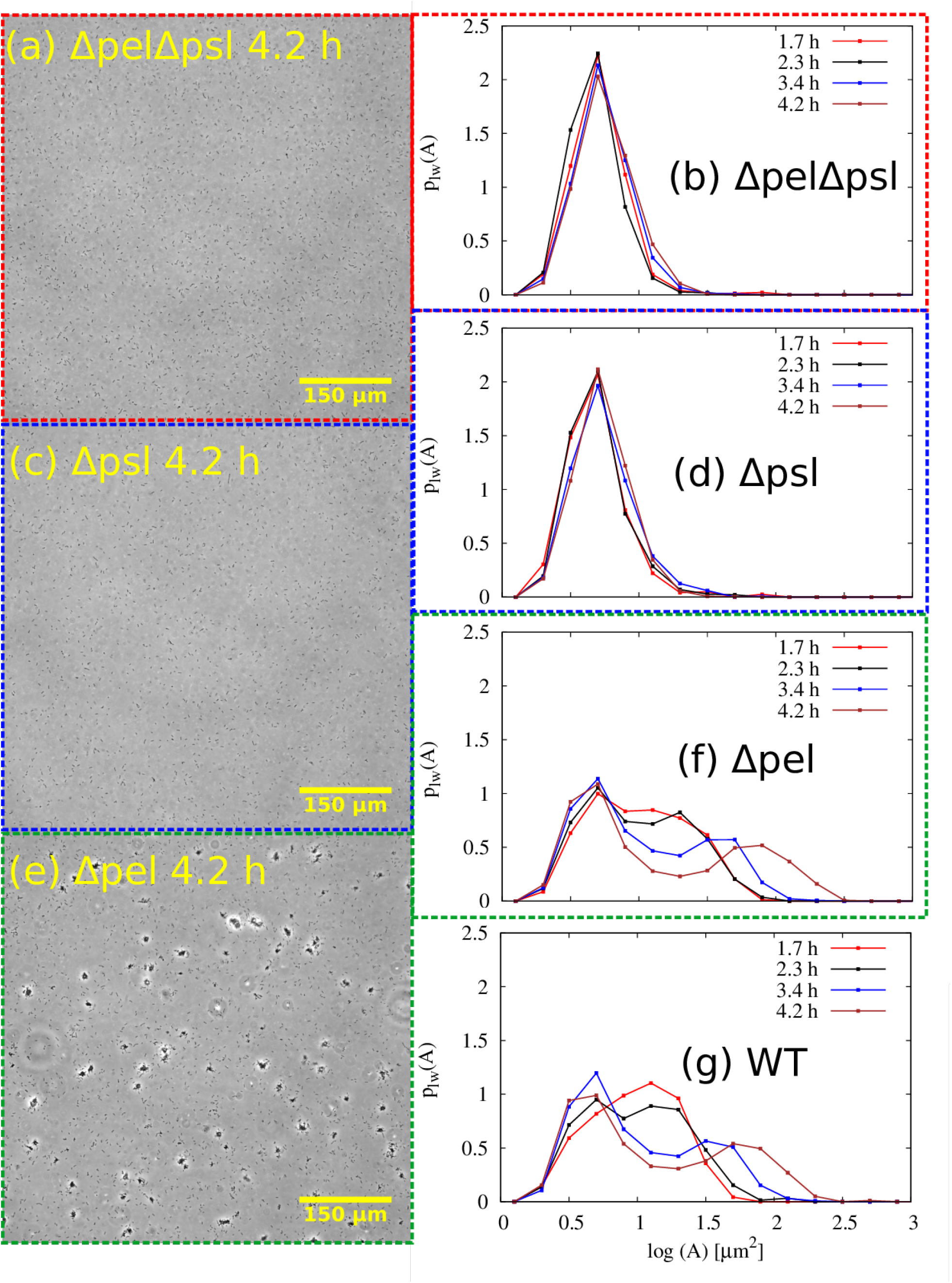
The Psl polysaccharide is required for aggregation. Representative images and distributions of aggregate sizes for the Δ*pel*Δ*psl*, Δ*psl*, Δ*pel* strains. (a), (c), and (e) Representative snapshots of the polymer mutants at 4.2 h. (b), (d), and (f) Logged and weighted aggregate size distributions at various times. For comparison, the distribution for the WT is also shown (g).

The Δ*pel* strain, however, did aggregate (Fig. 4(e)). Furthermore, quantification of the weighted aggregate size distribution *p*_*l w*_ (*A*) (Fig. 4(f)) showed results that were essentially identical to the wild-type (Fig. 4(g)). This suggests that Pel does not play a role in aggregate formation under these conditions.

To confirm that Psl is responsible for aggregate cohesion, we added the enzyme PslG, which degrades Psl (39, 40), to aggregated samples of the wild-type PAO1 strain. Addition of PslG (see Methods) results in loss of aggregates, as assessed both by microscopy (Fig. 5(a) and (b)) and by quantification of the weighted aggregate size distribution (Fig. 5(c)), for which the aggregate peak almost completely disappears in the presence of PslG. Since aggregates disappear when Psl is removed by the action of PslG, it appears that Psl provides a physical “sticking” force that holds the aggregates together.

**FIG 5.**
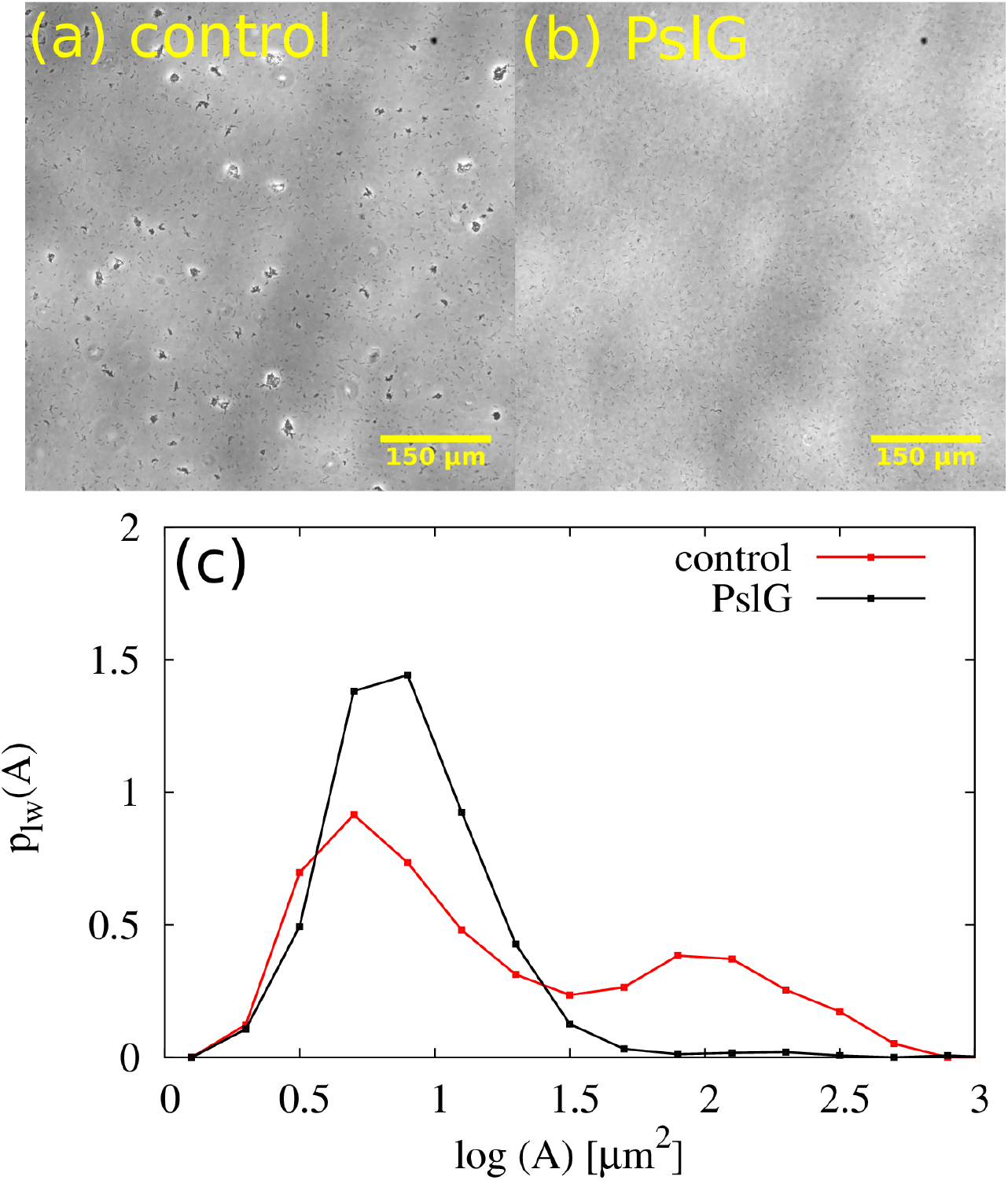
Psl is holding multicellular aggregates together. Representative phase contrast microscopy images of a control sample (a) and a sample treated with the glycoside hy-drolase PslG (b). (c) Aggregate size distributions for control and PslG-treated samples. Samples were incubated for 3.7 h, PslG was added (see Methods), and samples imaged 45 min later. Note there was a 20 min lag between imaging the control (4.1 h) and the enzyme-treated (4.4 h) samples.

### eDNA plays no role in exponential phase aggregation

Previous work suggests that, in addition to Psl, eDNA can play a role in aggregation of *P. aeruginosa* PAO1 (albeit in later-stage cultures) (16, 7, 6, 17). To test the role of eDNA in our exponential phase aggregates, we added DNase I, which degrades DNA, to aggregated samples from exponential cultures. DNase I had no effect on the aggregates in our samples, as observed by microscopy (Fig. 6(a)). Furthermore, quantification of the weighted aggregate size distribution revealed no significant effect of DNase I treatment Fig. 6(b). This suggests that eDNA plays no role in the formation of exponential phase aggregates.

**FIG 6.**
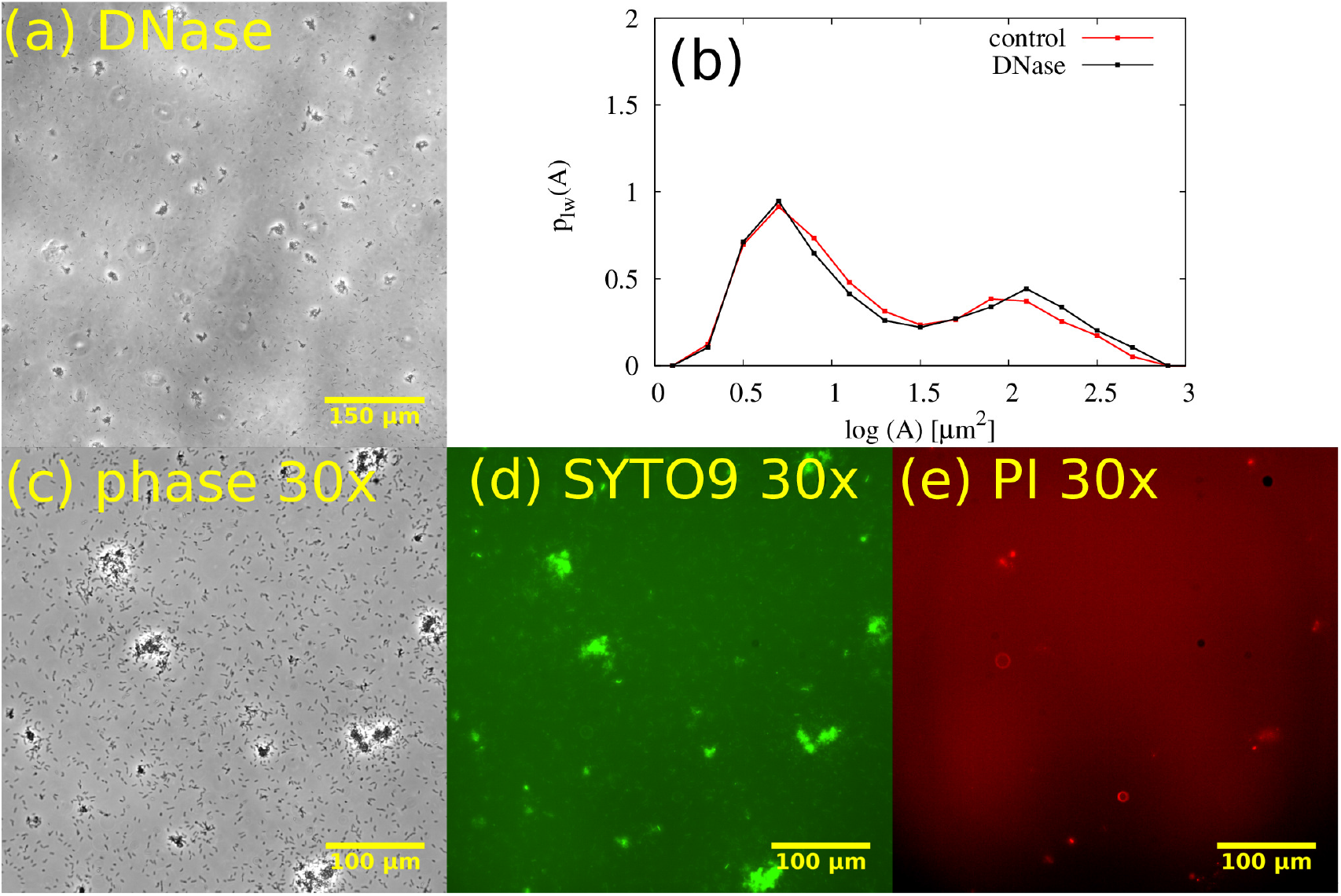
eDNA is not mediating aggregation. (a) Representative phase contrast image (4.1 h) of aggregated sample treated with DNase at 3.7 h after inoculation (see Methods). (b) Aggregate size distributions for control and DNase-treated samples. (c) - (e). Live-Dead staining of PAO1 cultures with SYTO9 (green) and PI (red) at 3.8 h after inoculation with dyes added 10 minutes before imaging (different data set to (a) and (b)). (c) 30× phase contrast image of representative snapshot shows aggregates and nonaggregated cells. (d) Corresponding image showing that many cells are stained with SYTO9 (green). (e) Corresponding image showing that few cells are stained with PI (red).

We also stained aggregated samples (Figs. 6(d) and (e)) from exponential phase cultures in our experiments with the dyes SYTO-9 (green) and propidium iodide (PI, red). SYTO-9 traverses the cell membrane and binds chromosomal DNA, so it can be used to locate cells in a fluorescence image. PI is a DNA-binding dye that does not traverse intact bacterial membranes, so it can be used to visualise eDNA, although dead cells will also appear as intense objects (16). We observed little signal in the red (PI) channel (Figs. 6(c) to (e)); contrast to stationary phase aggregates in Fig. 9), suggesting that exponential phase aggregates contain few dead cells, and little eDNA.

### Aggregation is mediated by cell-associated Psl

Our results imply that Psl mediates aggregation in exponential phase cultures of PAO1, with little or no role for either eDNA or Pel. However, Psl exists in two forms: a cell-free form that is secreted into the medium, and a cell-associated form that remains bound to the cell surface (36, 37).

To gain insight into the role of cell-free vs. cell-associated Psl, we reasoned that if cell-free Psl, in the medium, is important for cohesion, then diluting the culture should compromise the aggregates, since it will reduce the concentration of cell-free Psl that is present. In contrast, if aggregates are held together by cell-associated Psl they should remain intact after dilution of the culture, since the Psl is bound to the surfaces of the cells within the aggregate. Upon diluting aggregated samples from an exponential culture with fresh medium (in a 1:10 ratio), we observed, as expected, a decrease in the observed numbers both of aggregates and single cells (Fig. 7(a) and (b)). However, quantification of the weighted aggregate size distribution (Fig. 7(c)) revealed no difference in the size of the aggregates (Figs. 7(a) and (b)). This suggests that aggregate cohesion is mediated by cell-associated rather than cell-free Psl.

**FIG 7.**
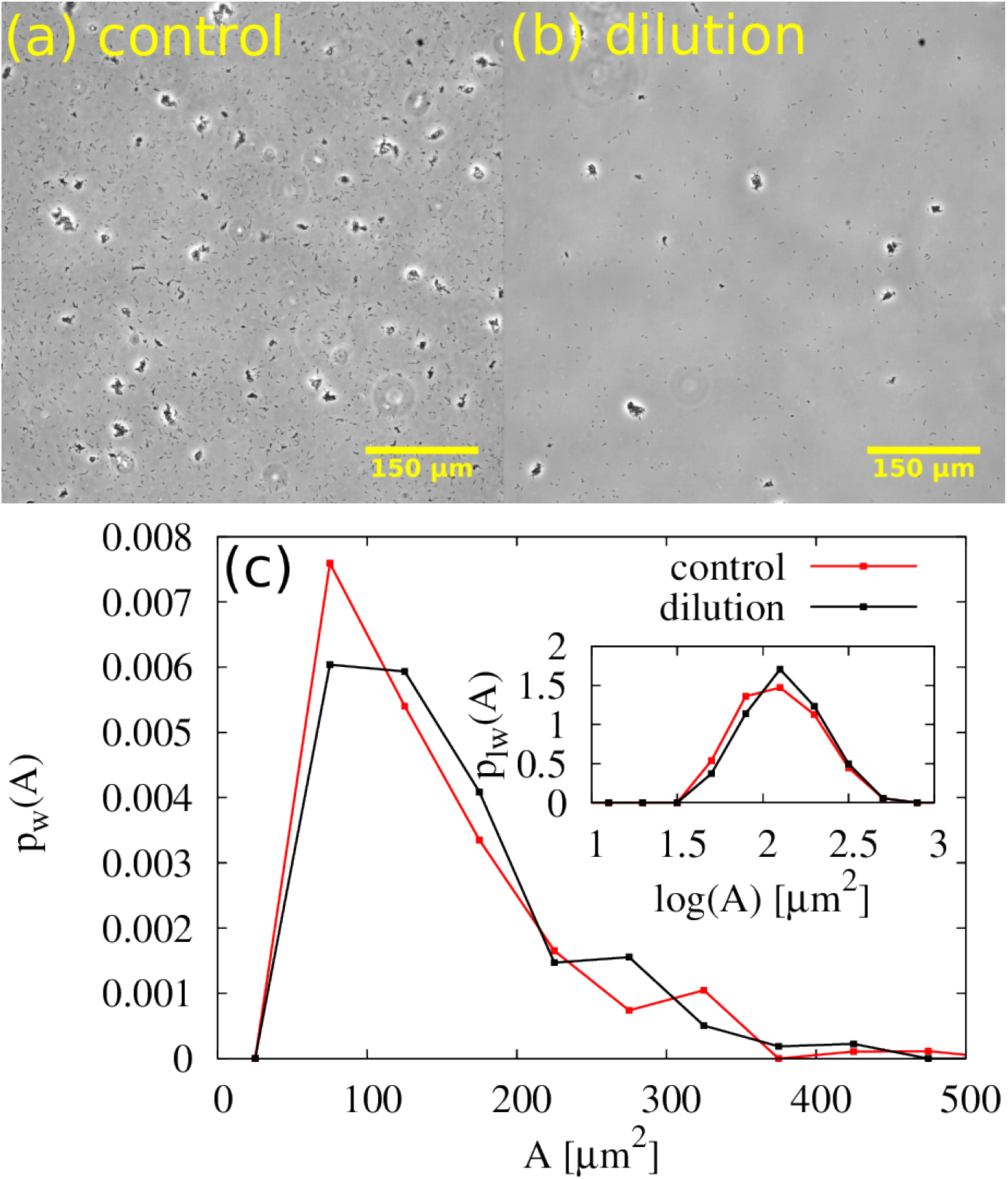
Aggregate size is unaffected by dilution. Upon dilution, the number of cells and aggregates in the eld of view is decreased but aggregate size is unaffected. (a) Representative phase contrast microscopy image of control sample of PAO1 at 3.6 h after inoculation. (b) Phase contrast image of diluted sample at 3.6 h after inoculation. (Note sample was incubated for 3.6 h. Then sample was diluted and imaged 5 minutes later). (c) Weighted distributions and log-weighted distributions (inset). Here, we are concerned with how dilution affects aggregate size, therefore entities less than 50 µm are not considered in the construction of the distributions.

### Aggregation mediated by cell-associated Psl is non-cheatable

To further test the hypothesis that cell-associated Psl mediates cohesion, and to understand its socio-evolutionary implications, we investigated the composition of aggregates in a mixed system of green Psl producers (Δ*pel*-GFP) and red nonproducers (Δ*pel*Δ*psl*-m-Cherry). In this system, the nonproducers might act as “cheaters”, since they could potentially join aggregates mediated by Psl-producers without paying the price of Psl production.

If aggregation were mediated by cell-free Psl, we would expect the Psl to be shared between producers and non-producers, since it is secreted into the medium. In this scenario, it would be likely that the non-producers could act as cheaters, benefiting from the secreted Psl. Therefore one would expect to see mixed aggregates, composed of both red and green cells (Fig. 8(a)).

**FIG 8.**
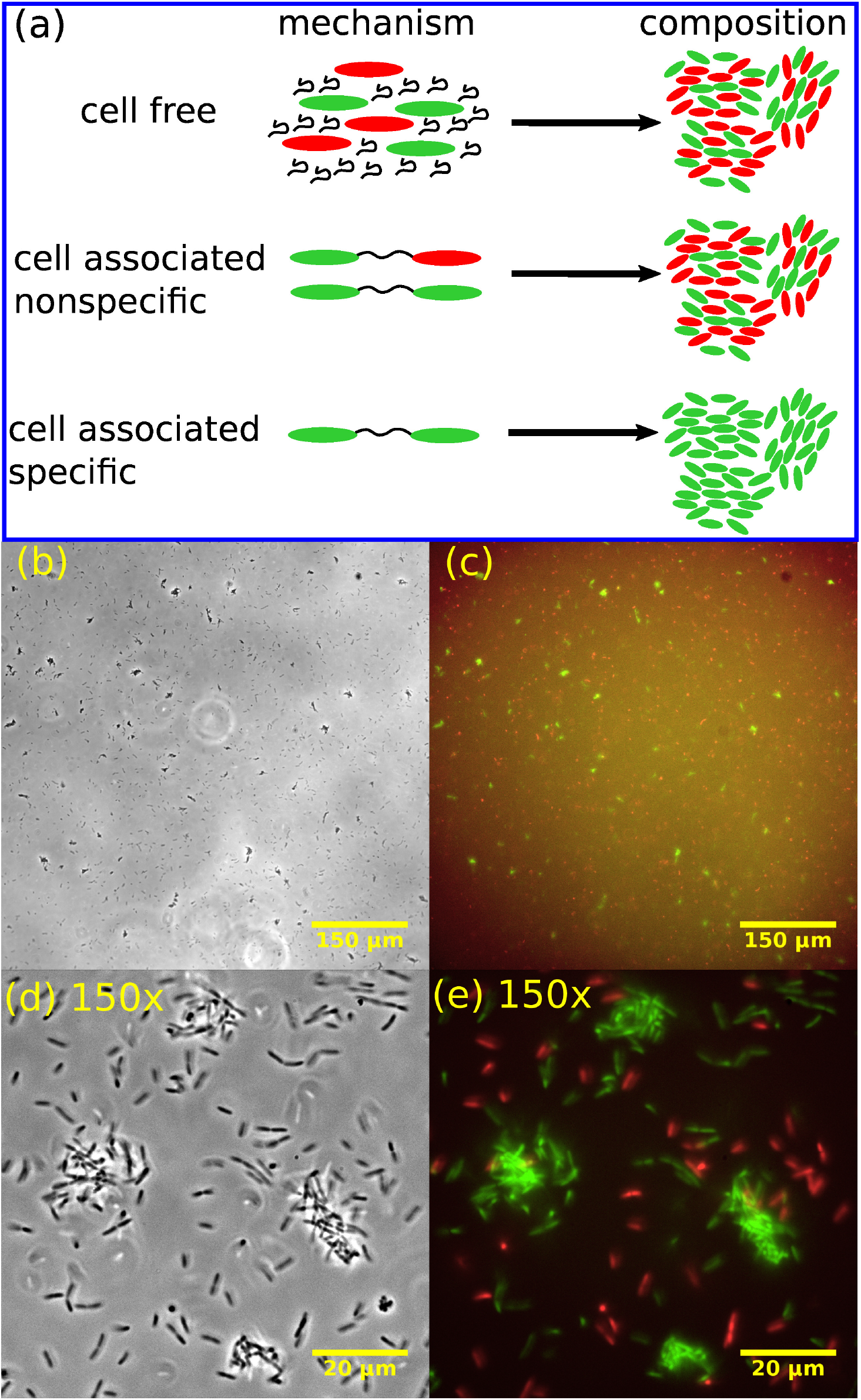
Cell-associated Psl mediates cell-cell cohesion. (a) Schematic outlining the composition of aggregates (right) resulting from the corresponding mechanism of Psl-mediated cohesion (left). (b) Representative phase contrast microscopy image (20× magni cation) taken 3.8 h after inoculation. (c) Superposed fluorescence images from green (Psl-producer, Δ*pel*-GFP) and red (nonproducer, Δ*pel*Δ*psl*-mCherry) channels corresponding to (b). (d) Representative phase contrast microscopy image (150× magnication) 4.0 h after inoculation. (e) Superposed fluorescence images from green (Psl-producer, Δ*pel*-GFP) and red (nonproducer, Δ*pel*Δ*psl*-mCherry) channels corresponding to (d).

If, on the other hand, the cell-associated form mediates aggregation, then we might expect either mixed aggregates or purely green aggregates. Mixed aggregates are expected if cell-associated Psl is “cheatable”, i.e. if it binds non-specifically such that it can form a “bridge” between a producer and non-producer. Purely green aggregates are expected if cell-associated Psl is non-cheatable, i.e. if it is specific in its binding, forming bridges only between producer cells (Fig. 8(a)).

Fluorescence microscopy of the red-green mixed culture revealed that aggregates consisted only of Psl-producing (green) cells (Figs. 8(b)-(d)). This confirms that, indeed, cohesion is mediated by cell-associated Psl, and furthermore, the interaction is noncheatable: cell-associated Psl bound to a producer cell leads to a cohesive interaction only with other producer cells, not with non-producer cells.

Previous research has shown that Psl can bind specifically to the matrix protein CdrA (52, 32) and so we speculated that the “specific” cell-associated Psl interaction that we observe might involve CdrA. However, we observed no discernible difference in aggregation for a mutant deficient in CdrA (Fig. S1).

### In stationary phase cultures, both eDNA and Psl mediate aggregation

Previous work has indicated a role for eDNA, or eDNA and Psl, in aggregation of *P. aeruginosa* PAO1 in late-log or stationary phase cultures (16, 17, 6, 7). We also obtained microscopy images of samples from overnight, stationary phase cultures. Aggregates were clearly visible in these samples (Figs. 9(a) and (b)), and these aggregates are stained by both PI and TOTO, indicating the presence of eDNA and/or dead cells (Figs. 9(c) and (d)). (TOTO, like PI, is a DNA-binding dye that does not traverse intact bacterial membranes, so can be used to visualise eDNA, although dead cells will also appear as intense objects (16))

**FIG 9.**
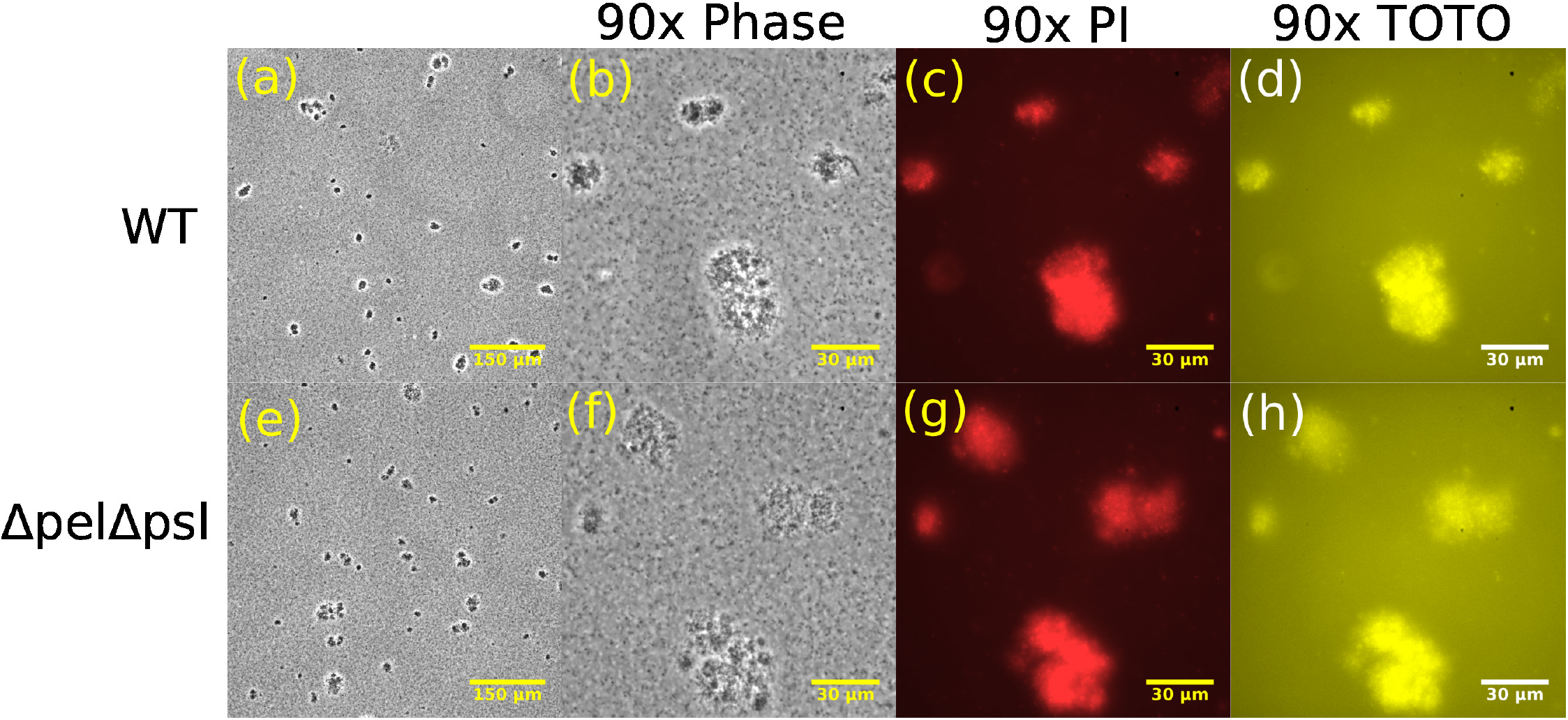
Aggregates of WT and Δ*pel* Δ*psl* strains are present in stationary phase culture. (a)-(d), Representative snapshots of WT aggregates in overnight culture. (a) Phase contrast 20× magni cation. (b) Phase contrast 90× magni cation. (c) Red luorescence channel, corresponding to image in (b), highlighting PI staining to extracellular DNA or intracellular DNA if cell membrane is compromised. (d) Yellow luorescence channel, corresponding to image in (b), highlighting TOTO staining to extracellular DNA or intracellular DNA if cell membrane is compromised. (e)-(f), Δ*pel*Δ*psl* snapshots corresponding to (a)-(d).

To confirm the role of eDNA and/or Psl in these stationary phase aggregates, we treated aggregated samples with DNase I (to degrade eDNA) or PslG (to degrade Psl), or both. Treatment with either DNase I or PslG resulted in smaller and less easily visible aggregates (compare Figs. 10(a),(b), and (c)). Simultaneous treatment with both enzymes led to complete loss of the aggregates. Therefore both eDNA and Psl are implicated in stationary phase aggregate formation in the PAO1 strain.

**FIG 10.**
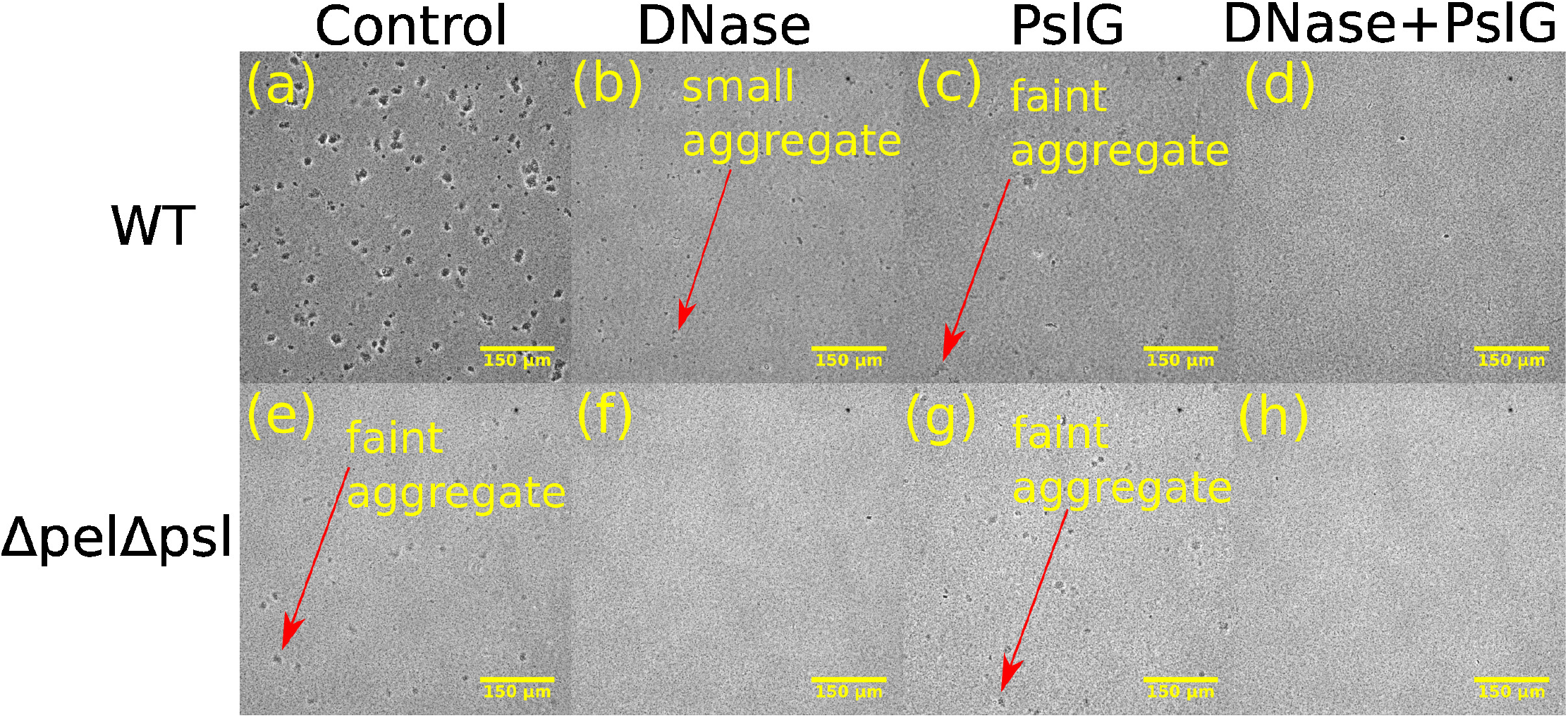
Both Psl and eDNA mediate aggregation in overnight stationary phase cultures Top row: WT strain subjected to treatment with DNase and PslG. Bottom row: Δ*pel*Δ*psl* strain subjected to treatment with DNase and PslG.(20x magnification)

Interestingly, the double polymer knockout strain Δ*pel*Δ*psl*, which cannot produce either Psl or Pel, also forms aggregates in overnight stationary phase culture, although these have a somewhat different appearance to those of the wild-type, appearing fainter in phase contrast images (Figs. 9(e) and (f); Fig. 10 (e)), similar to the wild-type aggregates after treatment with PslG (compare Fig. 10(e) to Fig.10(c)). The stationary phase Δ*pel*Δ*psl* aggregates stain with PI and TOTO (Fig. 9(g) and (h)), suggesting that their cohesion is provided by eDNA. Indeed, treating these aggregates with DNase I led to their complete disappearance (Figs. 10(f) and (h)), while treating them with PslG had no effect (Fig. 10(g)).

### Stationary phase aggregates do not seed exponential phase aggregates

Kragh et al. hypothesized a “snowball” mechanism, in which pre-existing small aggregates, present in the culture inoculum, could act as seeds for aggregate growth (7). Since our exponential phase cultures are inoculated from overnight stationary phase inocula that contain DNA and Psl-mediated aggregates (Fig. 9), we wondered whether these might act as seeds for the Psl-mediated aggregates that we observe in the exponential phase. To test this, we inoculated exponential cultures with stationary phase inocula that had been pre-treated by mechanical dispersion using a syringe, breaking up all aggregates (see Methods and Figs. S2(a) to (c)). Dispersal of the stationary phase aggregates in the inoculum had no effect on our results (Figs. S2(d) and (e)) suggesting that the aggregates which we observe in exponential phase are not seeded from pre-existing aggregates in the stationary phase inoculum.

## DISCUSSION

In this work, we investigated quantitatively the formation of suspended aggregates in exponentially growing liquid cultures of *P. aeruginosa* PAO1, compared to overnight stationary-phase cultures. We observe distinct types of aggregates for different growth stages. In exponential cultures, aggregation is mediated by Psl alone, aggregates appear *de novo*, and grow via a mixture of collisions and clonal growth. Psl in these aggregates is “non-cheatable” since Psl non-producers do not aggregate with producers. In contrast, in overnight stationary phase cultures both Psl and eDNA are involved in aggregation, although eDNA alone is suffcient.

Our results are consistent with previous observations that both eDNA and Psl are important for aggregation in overnight stationary phase aggregates (6, 7, 16, 17). Our finding that Psl mediates aggregation in exponential phase cultures is also consistent with previous work (7), however here we also show that it is cell-associated rather than cell-free Psl that plays the key role, with significant socio-evolutionary implications. Although previous studies have differentiated the roles of cell-associated and cell-free Psl in biofilm formation (25, 38, 39, 40), to the best of our knowledge, our study is the first to differentiate between the two forms in planktonic aggregates.

The extent to which the physiological state of bacteria in liquid-phase aggregates mirrors the surface-attached biofilm state has been a topic of discussion (6, 17, 7). In biofilms of *P. aeruginosa* PAO1, Psl has been found to play a role in initial attachment to surfaces (35, 53, 54, 34), microcolony and macrocolony formation (53), and is thought to be crucial in forming the matrix that holds cells together in more mature biofilms. eDNA, on the other hand, is mainly associated with late stage biofilm development (29), although it has also been found to be relevant for structural stability (34) and biofilm establishment (30, 31). For liquid-phase aggregates, our findings (and those of others (16, 17, 6, 7)) that Psl and eDNA play a role in the later stages, and only Psl at earlier times, suggest that there might be parallels between the development of biofilms and suspended aggregates.

In this work, we observed no role for Pel in aggregation of PAO1 in liquid. However, for other strains, such as PA14 and the PAO1 Pel-overexpressing strain (Δ*wspF*Δ*psl*), Pel has been implicated in biofilm formation (37, 41), as well as in aggregate formation in cystic fibrosis sputum (55). Therefore, it would be interesting to investigate whether Pel is important for planktonic aggregation in these strains. Previous work using atomic force microscopy suggests that Pel mediates cell-cell cohesion over a shorter range than Psl (35). Therefore we might expect Pel-mediated aggregation to proceed rather differently to the Psl-mediated aggregation studied here.

Interestingly, and in common with other authors (17, 7), we observe a population of non-aggregated cells that coexist with multicellular aggregates, even at late times (Figs. 1(c) and (d)). This seems somewhat surprising, since naively, we might expect that non-aggregated cells would eventually attach to aggregates. In future work, it would be interesting to investigate the origin of this coexistence. One explanation might be that the non-aggregated cells are phenotypically different from those in aggregates, for example, they produce less Psl. This in turn might be connected to the positive feedback regulatory circuit found in *P. aeruginosa* biofilms where Psl production stimulates increasing c-di-GMP which in turn stimulates increased Psl production (56), potentially leading to bistability in Psl production. One might also ask interesting general questions, such as how aggregation proceeds in suspensions in which cell-cell cohesion (“stickiness”) depends on the local cell density (57).

An innovative aspect of the work presented here is that we quantify the distribution of aggregate sizes. This allows us to determine, for example, that aggregates are indistinguishable for Psl-producers and non-producers. Quantification of the aggregate size distribution is also an important step towards the development of mathematical and computational models for bacterial aggregation (58, 59, 60, 61, 57, 62, 63, 64), similar to those that have been established for many years in the field of colloid science (65, 66, 67, 68, 69, 70, 71, 72). In particular, it is well established in the field of colloidal-polymer suspensions that micron-sized colloidal particles can aggregate either by depletion interactions, in which free-floating polymers in the suspension exert osmotic forces that push the colloids together (73, 74), or by polymer bridging interactions, where polymer molecules directly bind to the surfaces of the colloidal particles, sticking them together (75, 76). Depletion interactions have been implicated in several cases for bacterial aggregation in liquid suspension (77, 78, 79) and for host-polymer bacterial interactions in chronic infections (80, 81, 82) as well as experimental (83) and simulated (84) biofilms. In contrast, here we explicitly show that depletion is not relevant in exponential phase aggregation of *P. aeruginosa* PAO1, since the aggregates remain unchanged when we dilute the culture. Instead, we show that cell-associated Psl is responsible, suggesting a direct bridging interaction. Aside from the study by Jenning *et al* (55) which suggested the involvement of a polymer bridging interaction in CF sputum aggregates, to the best of our knowledge, our study is the first to point to a polymer bridging interaction in the context of planktonic aggregates. It will be important in the future to understand the molecular mechanisms behind this bridging, and subsequently construct appropriate models that can predict, for example, the time taken for a culture to aggregate.

**TABLE 1:**
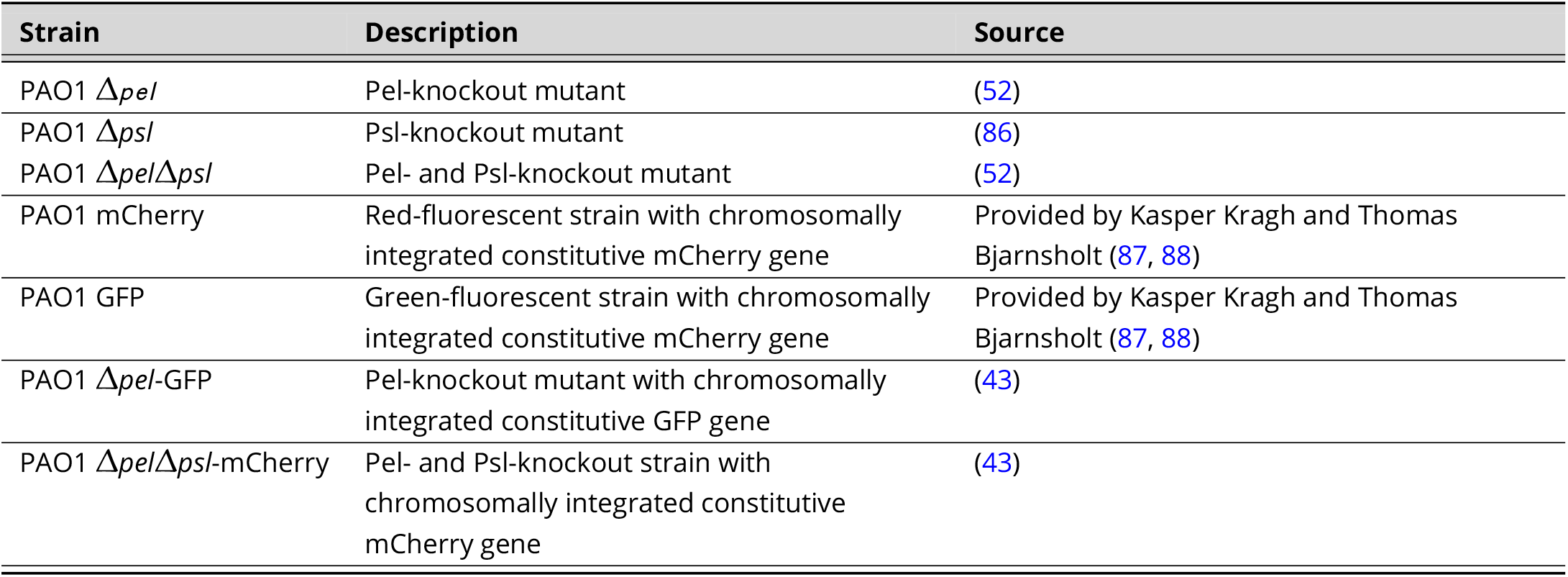
Strains used in this study

| Strain | Description | Source |
| --- | --- | --- |
| PAO1 $\Delta pel$ | Pel-knockout mutant | (52) |
| PAO1 $\Delta psl$ | Psl-knockout mutant | (86) |
| PAO1 $\Delta pel/\Delta psl$ | Pel- and Psl-knockout mutant | (52) |
| PAO1 mCherry | Red-fluorescent strain with chromosomally integrated constitutive mCherry gene | Provided by Kasper Kragh and Thomas Bjarnsholt (87, 88) |
| PAO1 GFP | Green-fluorescent strain with chromosomally integrated constitutive mCherry gene | Provided by Kasper Kragh and Thomas Bjarnsholt (87, 88) |
| PAO1 $\Delta pel$-GFP | Pel-knockout mutant with chromosomally integrated constitutive GFP gene | (43) |
| PAO1 $\Delta pel/\Delta psl$-mCherry | Pel- and Psl-knockout strain with chromosomally integrated constitutive mCherry gene | (43) |

Although quantitative measurements of the aggregate size distribution are very valuable, they are challenging in several ways. Our distribution *p* (*A*) is in fact the probability of observing an aggregate of area *A* on the bottom surface of a capillary (containing the liquid suspension) when viewed from beneath with an inverted microscope. Similar to previous studies (50), we use the area of the aggregates projected onto the surface as a proxy for aggregate size. Although this is not a true aggregate size distribution, owing to the fact the aggregates are 3-dimensional entities, it nevertheless allows one to compare aggregates at different stages during growth and also to compare aggregates between different strains. Combining this type of analysis with high throughput methods, like those used to assess coaggregation in aggregates of oral bacteria (85), will allow for a better understanding of aggregation in *P. aeruginosa* and in other strains that are clinically and industrially relevant.

The findings presented here highlight the fact that the physical, and perhaps also physiological properties of bacteria in planktonic aggregates might be very different depending on the environment in which the aggregates formed for example a stationary phase culture vs an exponential culture. Future work should investigate how the physical and biological properties of planktonic aggregates depend on their formation pathways, and the possible consequences of this for seeding of surface-attached biofilms (9, 10), as well as for antibiotic susceptibility in clinical infections, control of aggregation pathways and, potentially, our understanding of the evolution of multicellularity more generally.

## MATERIALS AND METHODS

### Growing liquid cultures

*Pseudomonas aeruginosa* strains were grown overnight from frozen stock cultures (80°C, 80% overnight cultures in LB and 20% glycerol) in 5 mL of Miller Lysogeny Broth (LB) at 37°C and 180 rpm for up to 16 h. Fresh culture was inoculated with a 1:1000 dilution of the overnight culture (100 µL of the overnight culture into 100 mL of fresh LB in a 500 mL Erlenmeyer flask). This is referred to as time zero (*t* = 0) for our exponential phase growth experiments. The freshly inoculated culture was incubated at 37°C and shaken at 180 rpm in an orbital incubator (Stuart, UK). The growth of the culture was monitored by taking OD_600_ measurements using a Cary 100 UV-Vis spectrophotometer (Agilent Technologies, USA).

### Imaging aggregates formed in liquid cultures

Samples were loaded into borosilicate glass capillaries (Vitrocom, 0.4 × 8.0 × 50 mm, volume ∼ 180 µL). The chamber was sealed using petroleum jelly to prevent leakage, and then placed onto a fully automated inverted microscope (Nikon TE300 Eclipse, Hamamatsu ORCA-Flash 4.0 camera) for imaging using phase-contrast and fluorescence microscopy. Once placed under the microscope, snapshots were taken at multiple horizontal (xy) positions on the capillary floor. Three types of objectives were used: a Nikon 20 × /0.5 objective; a Nikon extra-long-working distance 60 × /0.7 objective; and a Nikon 100 × /1.3 oil objective. For the higher resolution images (100× magnification) a different procedure was used. Instead of using a capillary tube, a small enclosure was created by attaching a gene frame to a microscopy slide. 100 mL of samples were loaded into this enclosure, which was then sealed with a cover slip, inverted, and placed onto the microscope for imaging.

Imaging at magnifications 30×, 90× and 150× was achieved using the microscope’s in-built 1.5× magnifier on the 20×, 60× and 100× objectives respectively. Pixel dimensions of 2048 × 2048 (1 × 1 binning) were used for imaging in phase contrast alone, whilst dimensions of 1024 × 1024 (2 × 2 binning) were used for combined fluorescence and phase contrast imaging.

### Syringing cultures

To assess the effect of aggregates in the initial inoculum, 1 mL sample of overnight culture was vigorously passed through a 20 G needle (BD, Spain) 5 times with a 1 mL syringe (BD, Spain). This was the control sample.

### Preparation and imaging of suspensions of PAO1 GFP and PAO1 m-Cherry strains

25 µL of overnight culture of PAO1 GFP (OD = 3.0) and 75 µL of overnight culture of PAO1 mCherry (OD = 1.0). Cultures were grown and imaged according to the methods outlined above. The dsRED channel (excitation FF01-554/23, emission FF01-609/54), and GFP channel (excitation FF01-474/27, emission FF01-525/45), were used for visualisation of the m-Cherry and GFP strains respectively.

### Computing the distribution of aggregate sizes

Phase contrast microscopy images (Fig. S3 left) were processed using FIJI (89) to generate the corresponding binary images (Fig. S3 right). First, a rolling ball background subtraction (FIJI, in-built function) was applied to correct for an uneven illuminated background. Here we used a rolling ball radius of 10 pixels. Binary images of dark object on light backgrounds were then generated with FIJI’s in-built threshold function using default and automatic settings. The resulting area of each dark object in each frame was then computed using FIJI’s analyse particles functions. This process was performed on many (∼ 10 − 40) microscopy images representing different fields of view (xy positions) in the sample.

In order to compute an area distribution, the outputted area sizes were used as input data for our in-house software (written in Fortran 90). Although it is somewhat arbitrary, we only considered aggregates with an area greater than 2 µm^2^. This is because visual inspection reveals that cells on the surface tend to have an area greater than 2 µm^2^ (Fig. S3; see also the individual cell peak in Fig 2 at log(*A*) ∼ 0.7 (10^0.7^ = 5 µm^2^)). The software then builds the distribution, *p* (*A*) by counting the number of aggregates *n*_*A*_ of area *A*.

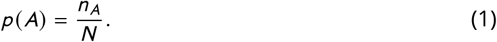

*N* is the total number of aggregates in the system and provides the normalisation

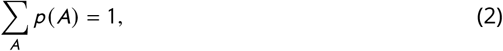

where the sum is over all aggregate sizes. We also computed a weighted distribution *p*_*w*_ (*A*) given by

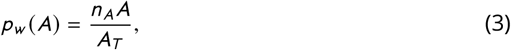

where *A*_*T*_ is the total area of aggregates in the system. Here we see that the number of aggregates *n*_*A*_ of size *A* is multiplied by *A* (weighted) to ensure that larger aggregates make a greater contribution to the distribution. The presence of *A*_*T*_ in the denominator ensures *p*_*w*_ (*A*) is normalised according to

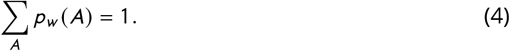

Given the large range of aggregate sizes observed in the system (2 − 1000 µm^2^), we also generated distributions in which the logarithm of the area (log_10_ (*A*)) was first computed before the frequency of the area is weighted by the area size. This has the effect of compressing the x-axis, thus making for easier visualisation.

Note that the black spot in the upper right corner of the microscopy images is a camera artefact and is subtracted from the binary images before computation of the aggregate size distributions.

### Enzyme treatment of exponentially growing cultures

For PslG treatment, 22 µL of 4.5 mg/mL of PslG solution (38) was added to a 20 mL universal flask along with 1 mL of liquid culture, which had been incubated for 3.7 h. The culture was then incubated for a further 40 min at 37°C while shaken at 180 rpm. For DNAse treatment, 100 µL of 100 µg/mL of DNase 1 (STEMCELL technologies, Canada) was added to 0.9 mL of liquid culture (incubated for 3.7 h) in a 20 mL universal flask. The culture was then incubated for a further 30 min at 37 degrees while shaken at 180 rpm. Samples were then visualised under the microscope. As a control, 1 mL of liquid culture (incubated for 3.7 h) was added to a 20 mL universal flask, without enzyme, and incubated for a further 30 min at 37°C while shaken at 180 rpm.

### Staining and luorescence imaging of exponentially growing cultures

For propidium iodide (PI)/Syto9 imaging, 20 µL of 100 µg/mL PI (Thermo Fisher) and 20 µL of 100 µM Syto9 (Thermo Fisher) were added to a 20 mL universal flask along with 2 mL of exponential phase culture, which had been incubated for 3.5 h. The sample was left for 15 min before being loaded into a capillary and imaged under the microscope. The dsRED channel (excitation FF01-554/23, emission FF01-609/54), and GFP channel (excitation FF01-474/27, emission FF01-525/45) were used for visualisation of the PI and SYTO9 dyes respectively.

### Dilution of exponential phase cultures

Exponential phase cultures were visualised under the microscope to check for the presence of aggregates in the bacterial suspension. After 3.5 h of growth, 120 µL of the exponential phase culture was added to 1080 µL of fresh LB media in a universal flask to give a dilution factor of 1:10. As a non-diluted control, 1200 µL of the exponential phase culture was added to a universal flask. Both samples were left to sit for 10 minutes before being visualised under the microscope.

### Preparation and imaging of suspensions of Δ*pel* GFP and Δ*pel*Δ*psl* m-Cherry

50 µL of overnight cultures of both Δ*pel* GFP (OD = 1.9) and Δ*pel*Δ*psl* (OD = 1.5) were added to 100 mL of fresh LB media to give a total dilution of 1:1000. The mixed culture was then incubated and images acquired as described above.

### Staining and luorescence imaging of stationary phase cultures

10 µL of 100 µg/mL Propidium Iodide (PI) (Thermo Fisher) plus 50 µL of 100 µM TOTO (Thermo Fisher) were added to 1 mL of stationary phase culture of WT cells (which had incubated for 22 h) in a 1 mL eppendorf tube. The sample was left for 15 minutes before loading a capillary and imaged under the microscope. The dsRED channel (excitation FF01-554/23, emission FF01-609/54), and YFP channel (excitation FF01-500/24, emission FF01-542/27) were used for visualisation of the PI and TOTO dyes respectively.

### Enzyme treatment of stationary phase cultures

30 µL of 4.5 mg/mL of PslG was added to 0.5 mL of stationary phase culture that had been incubated for 21 h. 100 µL of 100 µg/mL of DNase 1 (STEMCELL technologies, Canada) was added to 0.5 mL of stationary phase culture. As a control, 130 µl of PBS buffer was added to 0.5 mL of stationary phase culture. After addition of enzymes, samples were left for 30 minutes at room temperature before being loaded into capillaries and imaged under the microscope.

## ACKNOWLEDGMENTS

GM thanks the following funding bodies for financial support: EPSRC (Grant No. EP/ J007404); ERC (Grant No. 682237 “EVOSTRUC”); HFSP (Grant number RGY0081/2012); and NBIC/BBSRC/UKRI (Grant No.BB/R012415/1). RJA thanks the following funding bodies for financial support: EPSRC (Grant No. EP/J007404); ERC (Grant No. 682237 “EVOSTRUC”); HFSP (Grant number RGY0081/2012); NBIC/BBSRC/UKRI (Grant No.BB/ R012415/1); and the Deutsche Forschungsgemeinschaft (DFG) (Excellence Cluster Grant No. EXC 2051 - Project ID 390713860 “Balance of the Microverse”). PLH thanks the Canadian Institutes of Health Research (CIHR) (Grant Nos. 43998 and FDN154327). PLH was the recipient of a Tier I Canada Research Chair from 2006-2020. PB was supported in part by a Cystic Fibrosis Canada postdoctoral fellowship and a Banting Fellowship from CIHR. DJW and PH thank the National Institute of Health (NIH) (Grant Nos. R01AI077628, R01AI134895, and R01AI143916).

We thank Jochen Arlt, Thomas Bjarnsholt, Angela Dawson, Steve Diggle, Vernita Gordon, Yasuhiko Irie, Nick Jakubovics, Kasper Kragh, Alex Lips, Wilson Poon, Ross Slater and Sharareh Tavaddod for helpful discussions. We also thank Kasper Kragh and Thomas Bjarnsholt for providing strains.

For the purpose of open access, the author has applied a Creative Commons Attribution (CC BY) licence to any Author Accepted Manuscript version arising from this submission.

## SUPPLEMENTARY MATERIALS

**FIG 1.**
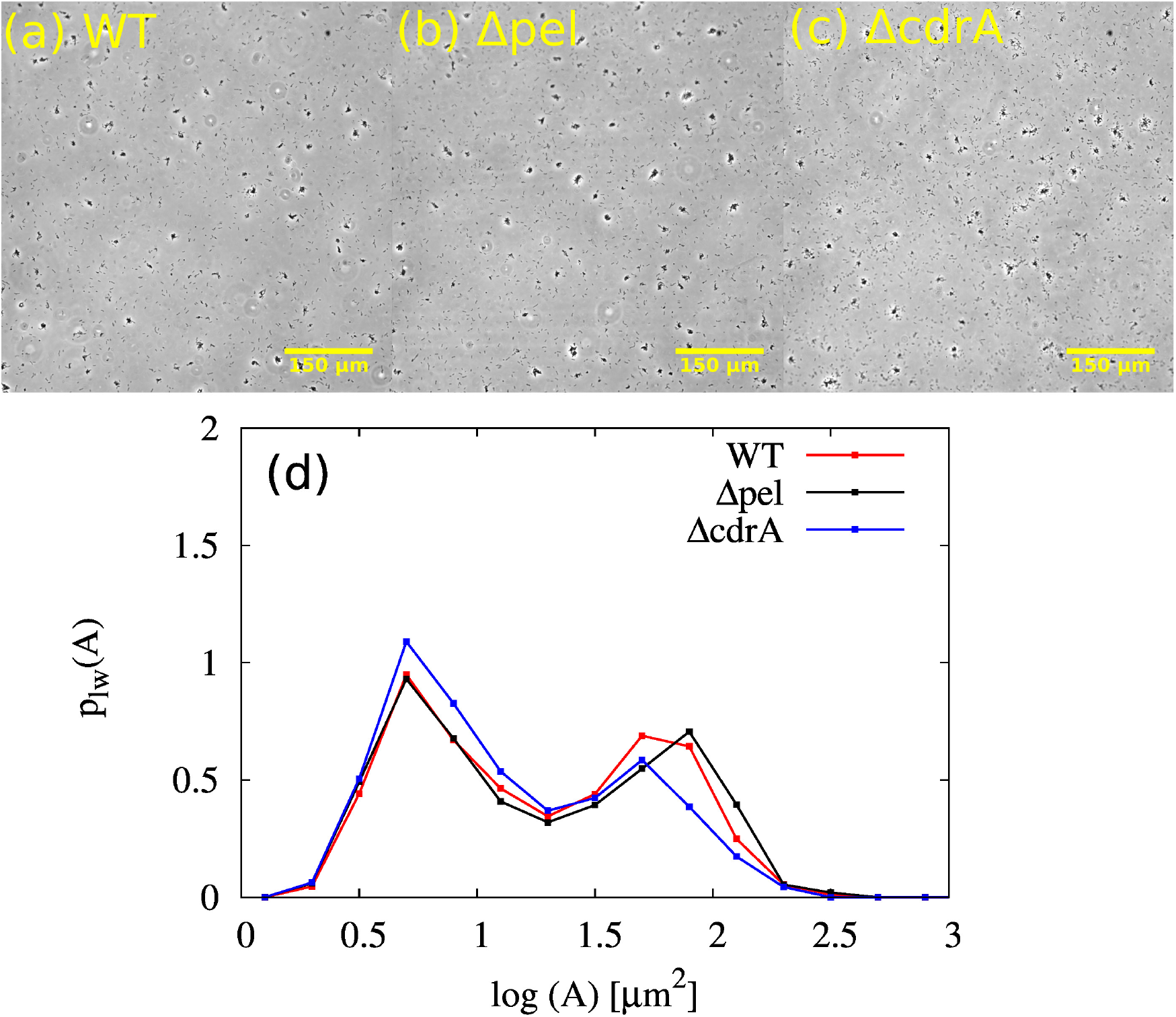
No obvious role of CdrA. Representative phase contrast microscopy images for (a) WT, (b) Δ*pel*, and (c) Δ*cdr A* at 2.5 h. (d) Corresponding aggregate size distributions

**FIG 2.**
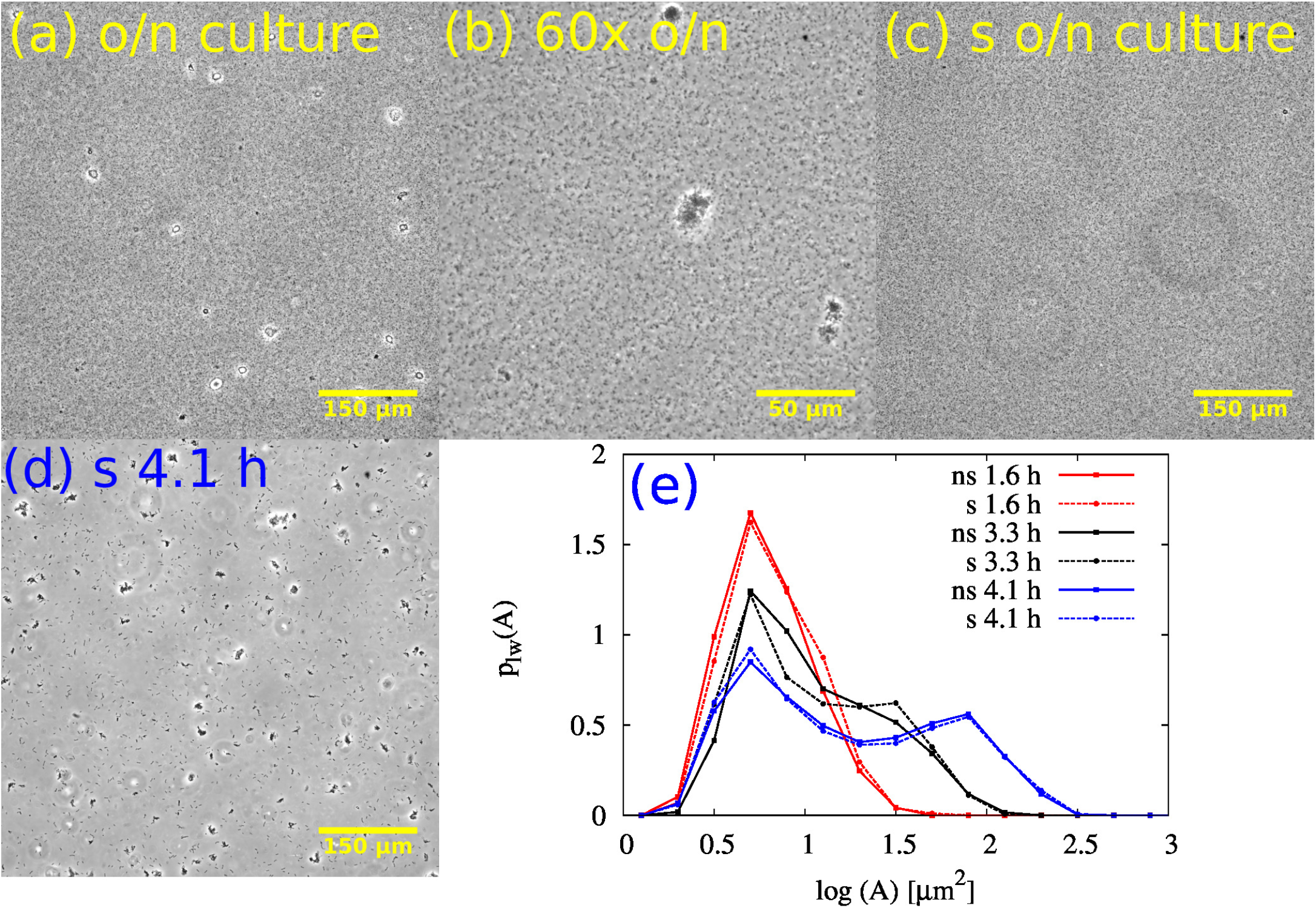
Exponential phase aggregates are not seeded by stationary-phase aggregates. (a) Phase contrast image showing aggregates in the overnight stationary phase culture. (b) 60× magni cation of overnight stationary phase culture in (a). (c) Phase contrast image of overnight stationary phase culture after mechanical disruption with syringe. (d) Phase contrast image of exponentially growing culture that was innoculated with a mechanically disrupted overnight culture. (e) Distribution of aggregate sizes in the exponential-phase cultures that were incoculated with (dashed lines) and without (solid lines) mechanically disrupted overnight cultures. s and ns denote syringed and nonsyringed inocula respectively.

**FIG 3.**
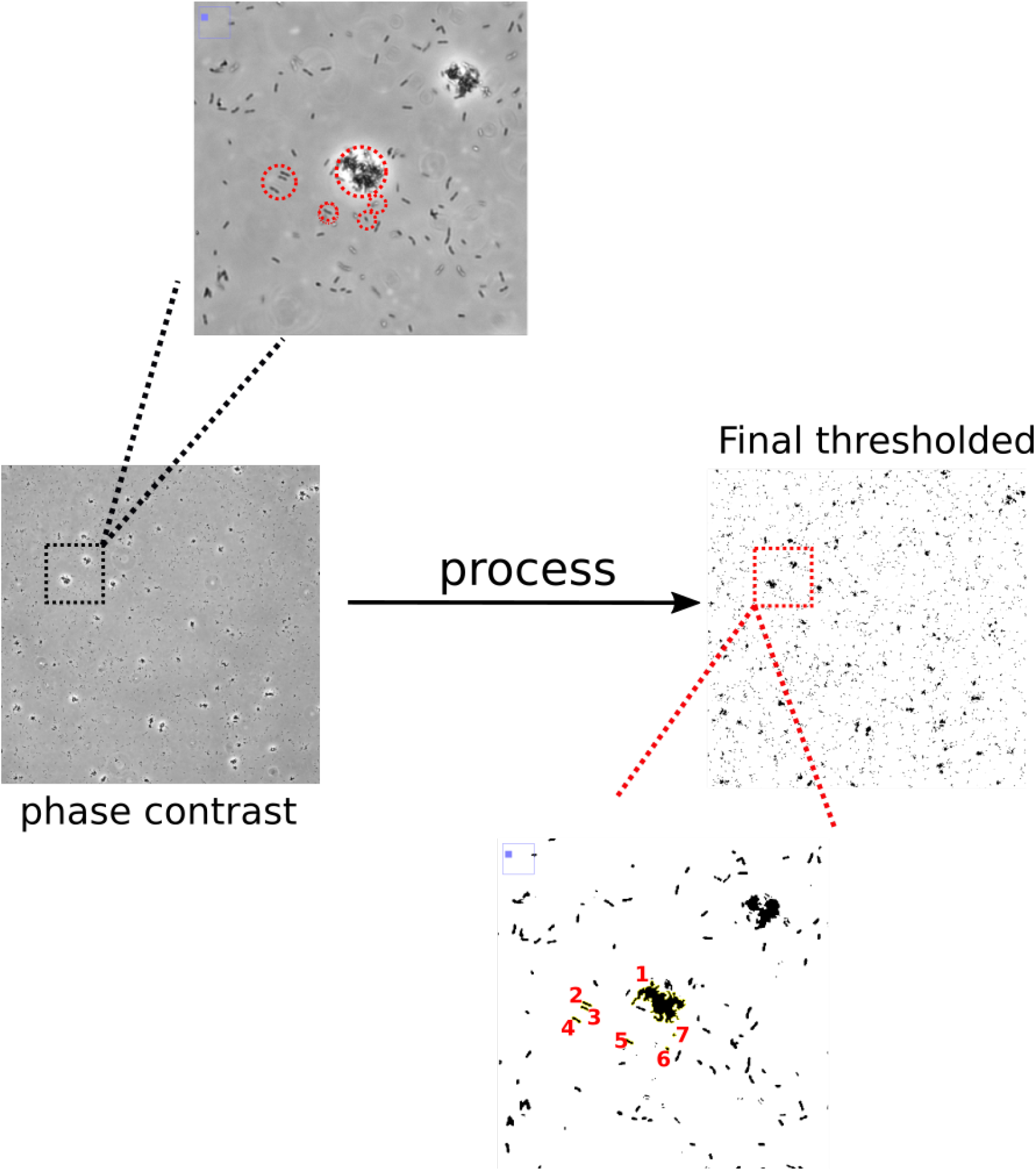
Computing the aggregate size distribution Left: Representative phase contrast microscopy image of an aggregated sample. Right: Corresponding thresholded image after processing. In the zoomed-in-region of the phase contrast image, the red circles highlight particular groups of unaggregated cells as well as aggregates. The sizes of the cells/aggregate (entities) within these red circles, numbered from 1 to 7 are: 180, 5.28, 4.12, 5.39, 4.65, 2.23, and 1.48 µm^2^ respectively

